# Phylogeny of the genus *Dolichopus* Latreille, 1796 and testing for phylogenetic signal in morphological characters and morphometric data

**DOI:** 10.1101/2022.02.24.481773

**Authors:** Mariya A. Chursina, Igor Ya. Grichanov

**Affiliations:** Natural Geographical Faculty, Voronezh State Pedagogical University, ul. Lenina 86, 394024 Voronezh, Russia; All-Russian Institute of Plant Protection, shosse Podbelskogo, 3, 196608, St. Petersburg – Pushkin, Russia

**Keywords:** Diptera, Dolichopodidae, *Dolichopus*, mitochondrial DNA, morphology, phylogeny

## Abstract

Four groups of characters were used to evaluate phylogenetic relationships among 50 species of the genus *Dolichopus*: 810 nucleotide characters (for cytochrome oxidase I), 18 continuous characters generated from a geometric morphometric analysis of wing shape, 12 relative characters of the leg morphometry and 42 traditional morphological characters. Subsequently, the common database was used to construct a phylogenetic genus tree and to study the presence of a phylogenetic signal in each group. In this study it was shown that the following characters have a significant phylogenetic signal: the thickening of the costal vein at the insertion point with *R*_*1*_, the color of the fore coxa, the presence of male secondary sexual characters on the forelegs, the color of the femora, the presence of long cilia on the hind femora, and the color of the lower postocular cilia. Traits of male wing shape, such us the relative position of the apex of *M*_*1+2*,_ the location of the posterior cossvein, the origin of the radial veins, and the position of the anal vein base, also showed a high phylogenetic signal. In addition, in males, the relative lengths of the first segment of the fore and mid tarsi showed a clear correlation with molecular data, which were interconnected with the presence of males leg modifications. In females, morphometric traits exhibited a less significant phylogenetic signal than in males, although, in most cases, the same traits evinced a high phylogenetic signal.

**Statements:** We (corresponding authors) certify that we have participated sufficiently in the conception and design of this work and the analysis of the data (wherever applicable), as well as the writing of the manuscript, to take public responsibility for it. We believe the manuscript represents valid work. We have reviewed the final version of the manuscript and approve it for publication. Neither has the manuscript nor one with substantially similar content under my authorship been published nor is being considered for publication elsewhere, except as described in an attachment. Furthermore, we attest that we will produce the data upon which the manuscript is based for examination by the editors or their assignees, if requested.

The present paper has not been submitted to another journal. All co-authors are aware of the present submission. All co-authors agree to the publication. The work was funded by RFBR and NSFC according to the research project No 20-54-53005.

---

The genus *Dolichopus* Latreille, 1796 currently includes more than 640 species (Grichanov, 2021) and is the largest genus of the Dolichopodidae family. Species of this genus are widespread and show the greatest diversity in the Palaearctic region. Phylogenetic analysis of the genus was carried out based on both morphological and molecular data (Zang & Yang, 2005; Bernasconi et al., 2007a, b; Germann et al., 2010, 2011); however, the studies included a limited number of species, concentrated mainly in the eastern part of the Palaearctic. In this study, for the first time, we have included in the analysis the molecular traits of 20 *Dolichopus* species, which, among other things, have central- and eastern Palaearctic habitats.

For the taxonomy of *Dolichopus* species the following morphological characters are traditionally used: color of eye and postocular setae, postpedicel shape and antenna color, chaetotaxy and color of legs, wing morphometric characters and characteristics of male hypopygium. In addition, various traits of male sexual dimorphism are widespread: elongated postpedicel, modified arista, modifications of several segments of the legs, changes in the color of the wings, and thickening of the costal vein. The advent of geometric morphometry methods made it possible to identify the shape of the wing as a new category of characters, which is now widely used both for phylogenetic, taxonomic and diagnostic purposes (Schutze et al., 2012; Pepinelli et al., 2013; Torres et al., 2015). The resulting wing shape variations can potentially contribute to a general phylogenetic hypothesis based on molecular data and traditional morphological characters. The presence of a phylogenetic signal in characters of a particular category is still under discussion (Bernasconi et al., 2007b; Chursina & Grichanov, 2019), since the emergence of similar structures can be explained not only by the evolutionary proximity of species, but also by convergent evolution, as described, for example, for characters of sexual dimorphism (Bonduriansky, 2006; Chursina, 2019).

A detailed analysis of the phylogenetic significance of various groups of characters is important for understanding evolutionary trends in the family. Therefore, the purpose of this study was not only to analyze to analyse the related relationships between *Dolichopus* species, but also to assess the phylogenetic signal in characters of three main groups: traditional morphological characters (including chaetotaxy characters, color characteristics, and the characters of sexual dimorphism), legs morphometric characters and wing shape.

## Materials and methods

In this study, an effort was made to sample as widely as possible from the *Dolichopus* species. The general data matrix included characters of 50 Dolichopus species, including molecular data of 20 species were examined for the first time. In the course of the study, molecular methods, methods of traditional morphology, as well as methods of geometric morphometry were used.

### Samples

We studied a total 2195 specimens from 50 *Dolichopus* species from the entomology collection of the Voronezh state university (Voronezh, Russia) and the personal collection of authors (Table 1). Since it was previously shown that among the species of the subfamily there is clear sexual dimorphism both in the wing shape (Chursina & Negrobov, 2018) and in the legs morphometric characteristics (Chursina & Negrobov, 2020), males and females require separate consideration.

### Molecular data analysis

The analyzed molecular matrix included molecular sequences of the mitochondrial gene encoding cytochrome c oxidase (COI) (810 characters). The study included both sequences previously deposited in GenBank (GenBank, 2021) and sequences carried out especially for this study by the Sintol Enterprise (Russia). In total molecular sequences of 50 species was studies. Amplification and sequencing were performed using the methods and primers described in previous studies (Simon et al., 1994; Simmons & Weller, 2001; Bernasconi et al., 2007a, b). The sequences were aligned manually using BioEdit multiple alignment software (Hall, 1999).

Phylogenetic reconstruction was carried out using four methods: the maximum parsimony method (MP), the maximum likelihood method (ML), the minimum evolution method (ME), and the neighbour-joining method (NJ) in MEGA software (Kumar et al., 2018). Reliability of inner branches was estimated by the bootstrap method based on 1000 pseudoreplicates.

### Morphometric analysis of legs

Measurements were made by the photos of the specimens using the ImageJ software (1.53b) (Schindelin et al., 2015). Nine morphometric characters of the legs were measured: the lengths of the fore-, mid-, and hind femora (F1, F2, and F3), the fore-, mid-, and hind tibia (T1, T2, and T3), and the first segment of the fore-, mid-, and hind-tarsi (tar1, tar2, and tar3). Then the following twelve relative signs were calculated: the ratio of the lengths F1 to T1, F1 to tar1, F1 to F2, F1 to F3, T1 to tar1, T1 to T2, T1 to T3, F2 to T2, F2 to tar2, T2 to tar2, F3 to T3, and F3 to tar3. To examine relationship among species based on legs morphometric characters, unweighted pair group method with arithmetic average (UPGMA) was performed using PAST 3.09 software (Hammer, 2001).

### Geometric morphometric analysis of wing shape

To describe wing shape nine homologous Type 1 landmarks, placed in the vein junctions and vein terminations, were used (fig. 1). Each landmark have been digitized using TpsDig-2.32 software (Rohlf, 2008).

**Fig. 1.**
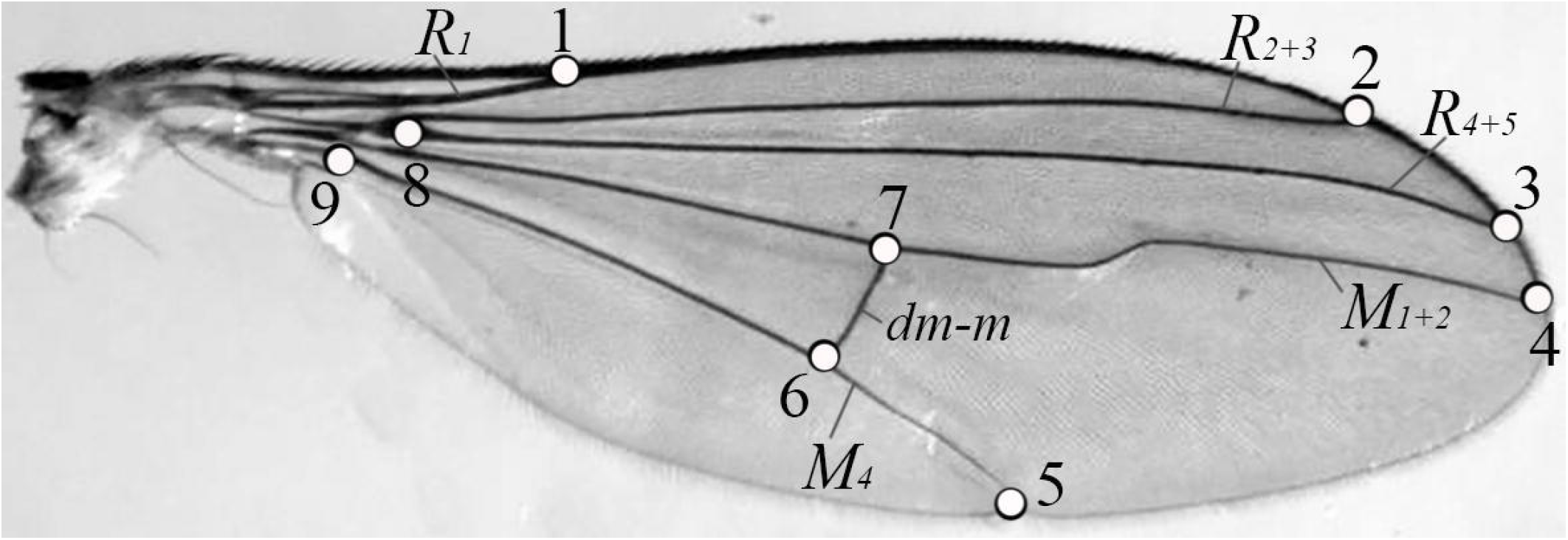
Positions of the landmarks used for studying wing shape (*Dolichopus latilimbatus*, male).

**Fig. 2.**
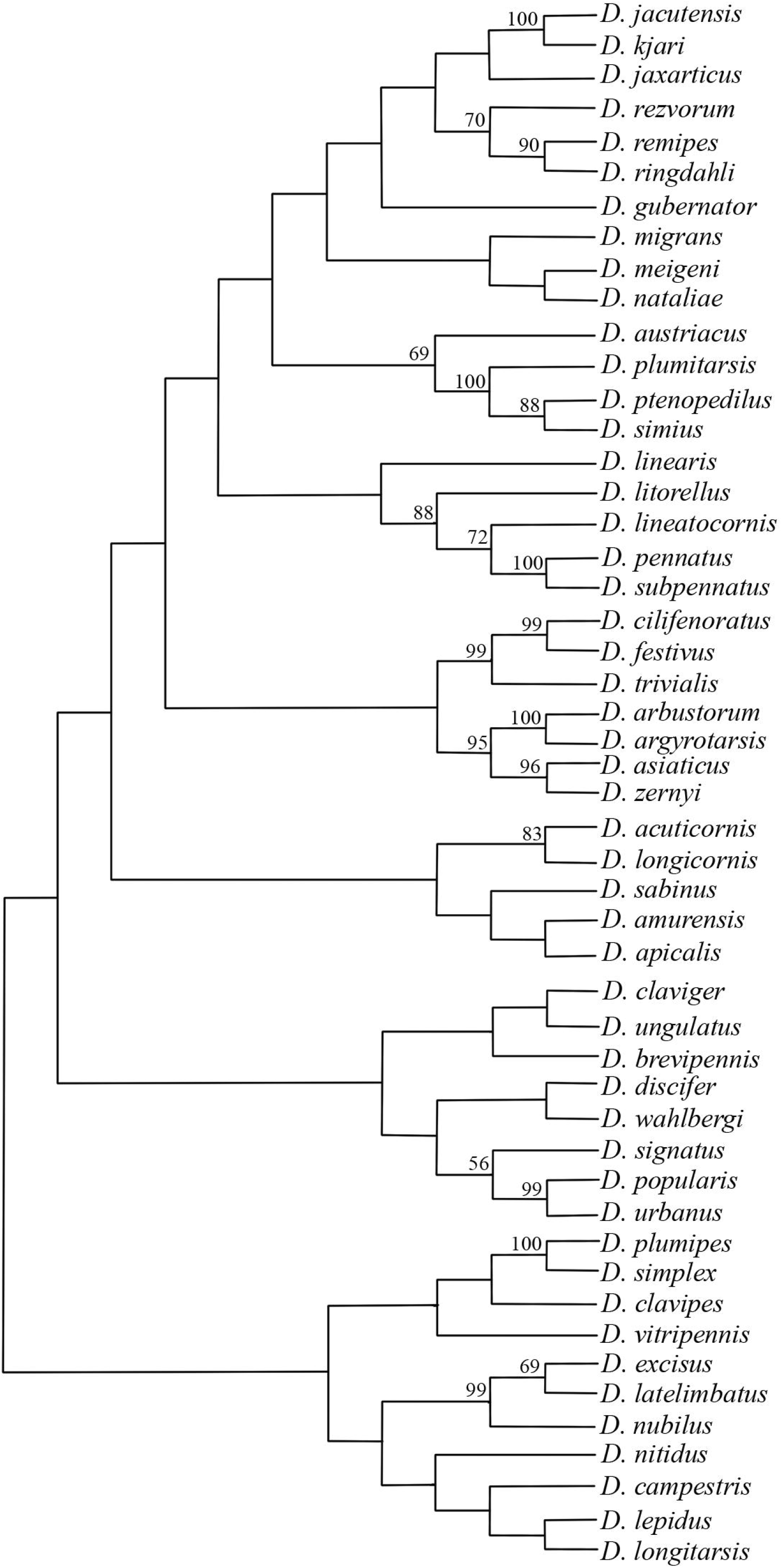
Maximum likelihood tree from 100 bootstrap replicates, obtained from molecular data (-Ln = 12732.87). Bootstrap support values higher than 50% are indicated above branches.

Centroid size (square root of the sum of squared distance between each landmark and the wing centroid) was used as the measure of size for each wing. Procrustes coordinates obtained from landmark data were used for further statistical analyses of wing shape. For this purpose, all wings were superimposed and differences in the position of the landmark points were then examined. The superimposition was done with the Procrustes technique (wings are scaled to a unit centroid size, superimposed on an origin so that their mean x and y coordinates becomes (0, 0), the landmarks are finally rotated so that the distance between all the landmarks of all specimens and a consensus configuration becomes minimal) (Rohlf, 1999). Then, analysis was carried out using the methods of multivariate statistical analysis in software MorpholJ (Klingenberg, 2011) и Statistica for Windows (version 10). To examine relationship among species based on wings morphometric characters, unweighted pair group method with arithmetic average (UPGMA) was also performed using PAST 3.09 software (Hammer, 2001).

### Traditionally used diagnostic morphological characters

Forty-two morphological characters that have diagnostic value in identification keys have been selected for comparative analysis. Characters states were presented in matrix.

I. Head
  1. Yellow face (1). White face (0). Dark face (2).
  2. Frons metallic-green, shining (1). Frons dense pollinose (0).
  3. Antennae mainly black (1). Antennae entirely black (0). Antennae mainly yellow (2).
  4. Postpedicel elongated, at least 1.5 times longer than high (1). Postpedicel short, nearly as long as high (0).
  5. Arista strongly plumose (1). Arista bare or weakly pubescent (0).
  6. Arista with enlarged apicaly (1). Arista simple (0).
  7. Lower postocular bristles yellow (1). Lower postocular bristles black (0).
    II. Thorax
  8. Clutch of small setae in front of posterior spiracle is present (1). Clutch of small setae in front of posterior spiracle is absent (0).
  9. Dark spot above notopleuron is present (1). Dark spot above notopleuron is absent (0).
    III. Wings
  10. Males wing at least partly darkened (1). Males wing hyaline or weakly smoky (0).
  11. Costa thickened before the tip of first radial vein (R_1_) is present (1). Costa thickened before the tip of first radial vein (R_1_) is absent (0).
  12. Costa thickened at tip of first radial vein (R_1_) is present (1). Costa thickened at tip of first radial vein (R_1_) is absent (0).
  13. Anal lobe well developed (1). Anal lobe obtuse (0).
  14. *R*_*4+5*_ and *M*_*1+2*_ subparallel beyond convergent apically (1). *R*_*4+5*_ and *M*_*1+2*_ subparallel apically (0).
  15. Vein *M*_*1+2*_ beyond crossvein *dm-m* with strong curvation (1). Vein *M*_*1+2*_ beyond cross-vein dm-m with weak curvation (1).
  16. *M*_*2*_ is absent (1). *M*_*2*_ is present as stub (0).
  17. Apical part of M_4_ in 1–2 times longer than *dm-m* (1). Apical part of M_4_ more that two times longer than *dm-m* (0). Apical part of M_4_ approximately equal *dm-m* (2).
  18. Squamal fringe dark, although some light-colored hairs may be present (1). Squamal fringe entirely dark (0). Squamal fringe entirely pale (2).
  19. Halters yellow with dark spot (1). Halters entirely dark (0). Halters entirely yellow (2).
    IV. Legs
  20. Fore coxa yellow (1). Fore coxa dark (0).
  21. Fore femora mainly or entirely yellow (1). Fore femora mainly or entirely dark (0).
  22. Fore tibia mainly or entirely yellow (1). Fore tibia mainly or entirely dark (0).
  23. Male fore tibia with long apicoventral seta (1). Long apicoventral seta is absent (0).
  24. Some segments of fore tarsus with regular erect setae (1). Fore tarsi simple (0).
  25. Some segments of fore tarsus enlarged (1). Fore tarsi simple (0).
  26. Some segments of fore tarsus have a changed color (1). Fore tarsi simple (0).
  27. Mid femora with one apical bristle (0). Mid femora with two or more apical bristles (1).
  28. Some segments of mid tarsus with regular erect setae (1). Mid tarsi simple (0).
  29. Some segments of mid tarsus enlarged (1). Mid tarsi simple (0).
  30. Some segments of fore tarsus have a changed color (1). Mid tarsi simple (0).
  31. Fore coxa mainly or entirely (1). Fore coxa dark (0).
  32. Hind femora yellow, darkened apically (1). Hind femora entirely yellow (0). Hind femora mainly dark (2).
  33. Hind femora with one apical bristle (0). Hind femora with two or more apical bristles (1).
  34. Hind tibia mainly yellow (1). Hind tibia mainly dark (0). Hind tibia entirely yellow (2).
  35. Hind legs modified: hind tibia thickened of hind legs ornamented (1). Hind leg simple (0).
  36. Cilioratum is present, small (1). Cilioratum is absrent (0). Cilioratum is present, elongated (2).
  37. The first segment of hind tarsi with one strong dorsal bristles (1). The first segment of hind tarsi without dorsal bristles (0). The first segment of hind tarsi with two or more strong dorsal bristles (2).
  38. At least the first segment of hind tarsi yellow (1). Hind tarsi entirely dark.
  39. Hind femora with long ventral hairs (1). Hind femora with short ventral hairs (0).
    V. Males genitalia
  40. Apical epandrial lobe elongated and think (1). Apical epandrial lobe short and wide (0).
  41. The ratio of the length of epandrium and cerci is less than 2 (1). The ratio of the length of epandrium and cerci is more than 2 (0).
  42. Cerci well developed, subtriangular or или rhomboid, with strong apical bristles (1). Cerci rather small, oval or rounded (0).

The significance of the inner branching pattern was estimated by a bootstrap analysis with 1000 pseudo-replicates. As a measure of phylogenetic signal of legs morphometric characters, we used Pagel’s lambda (λ) (Pagel, 1999) and Blomberg K-statistic (Freckleton et al., 2002). A Pagel’s lambda is an indicator that can take a value from zero to one, and a value close to one indicates a more significant phylogenetic signal in character. To calculate Pagel’s lambda, the phylosyg function phytools package (Revell, 2012) was used in R environment (R Development Core Team, 2014). Blomberg K-statistic also takes values from zero to one, but if the phylogenetic signal is very high, then K-statistic can rise over one. To calculate Blomberg K-statistic, the Kkalk function picante package was used in R environment (Kembel et al., 2010). For testing purpose the indications of differences of the metric from 0, a p-value was obtained by randomizing the trait data 1000 times.

## Results

An overview of the identified species groups of species with the corresponding indices of bootstrap support is presented in the table 2. According to trees, obtained from molecular data, ten groups of species are most consistently distinguished.

1. The morphological similarity of the species *D. arbustorum, D. argyrotarsis, D. asiaticus* and *D. zernyi* shares traits such as legs colour (femora, fore coxa and tibia are yellow), and the characteristics of the wing, in which thick costal vein are absent and the postocular cilia is pale. Given that these traits are characteristic of a greater number of *Dolichopus* species, there is no evidence of this group of species, including those with morphometric characteristics of legs and wings, in these produced from morphological data. In addition, *D. argyrotarsis* males have modified mid tarsus: the third, fourth and fifth segments are silvery, while the mid tarsus of the other species in the group are plain. The wing shape, as well as the morphometric characteristics of the legs of *D. argyrotarsis*, bear striking similarity to *D. lineatocornis*, a species whose males also have a modified mid tarsus (Fig. 3, 4).
2. Males of *D. acuticornis* and *D. longicornis* have elongated postpedicel, costal thickening (but with different localization), similar leg colour features except for the hind legs, as well as similar characteristics of the cerci. In addition, the wing shapes of both species bear a marked similarity (Fig. 3).
3. *Dolichopus cilifemoratus* Macquart species group. One of the most reliably identified phylogenetic groups of species, including in other studies (Bernasconi et al., 2007b), is a group that includes *D. cilifemoratus, D. festivus* and *D. trivialis*. All three species possess significant similarities according to traditional morphological characters: yellow femora, fore coxa, and fore and mid tibiae, the presence of long hairs on the hind femur, but also in terms of formal wing characters: the costal vein is thickened at the tip of *R*_*1*_, the length of the apical segment of *M*_*4*_ is approximately equal to the length of the *dm-m*. In addition, males of all three species have a similar modified of the fore tarsus - all segments have short erect setae. However, there is no significant similarity in the morphometric characteristics of the wings and legs. Other researchers (Pollet et al. 2010) also include *D. arbustorum* in this group, which has a number of characters in common with the species of this group, but differs from all three by a hypandrium and basiventral epandrial lobes that are entirely symmetrical, costal vein lacks thickenings and of the minute erect setae on the fore tarsus. According to our data, *D. arbustorum* has no similarity either in molecular characteristics, or in the wing shape and legs morphometric characteristics.
4. The allocation of the *Dolichopus plumipes* species group (*D. plumipes* and *D. simplex*) is not confirmed by any unique combination of morphological characters among the studied species, as well as by the similarity of morphometric characters. An important diagnostic character of *D. plumipes* male is the bilaterally feathered first segment of the mid tarsus in combination with a thin mid tibia. According to the morphometric characters of the leg, *D. plumipes* bear striking similarity with *D. wahlbergi*, whose males have a similar modification of the mid legs (Fig. 4). *Dolichopus simplex* is remarkable in absence of any male secondary sexual characters. In the study by Pollet et al. (2010), the analysis of a mitochondrial DNA dataset, consisting of 1702 characters, made it possible to classify within the *Dolichopus plumipes* group the species *D. discifer* (=*D. nigricornis* Becker, 1917, nec Meigen, 1824), which is the only species in the group having modified fore tarsus (Khaghaninia et al., 2014). This result is interesting in that, if *D. discifer* is included in this group of species, it will be the only group identified based on the basis of molecular data, which includes both species with modified fore tarsi and species with modified mid legs.
5. The group *D. popularis* and *D. urbanus* identified in the molecular analysis, is the only case in our study where species with similar modification of male mid legs also showed a similarity based on molecular data, although in terms of morphological characters used to diagnose *Dolichopus* species, *D. popularis* is more similar to species such as *D. lineatocornis, D. pennatus* and *D. signatus*. In the morphometry-based tree, *D. popularis* is always clustered together with species that have male secondary sexual characters on the mid leg (*D. plumipes* or *D. subpennatus*).
6. The group *Dolichopus latilimbatus*, including *D. nubilus, D. latelimbatus* and *D. excisus*, was identified earlier (Bernasconi et al., 2007b) based on a number of morphological characters, of which its unique combination is particularly important to highlight, such as pubescence facial hair and a darkened apex of the hind femur.
7. The group *Dolichopus kjari*, including *D. kjari* and *D. jacutensis*, is characterized by combination of characters such as a white face and shiny frons, pale lower postocular cilia, yellow fore coxae, femora, entirely yellow fore tibia and entirely dark hind legs. Males of both species of the group have a thickening on the costal vein and long apicoventral setae on the fore tibia.
8. In the *Dolichopus subpennatus* species group (*D. pennatus* – *D. subpennatus* – *D. lineatocornis* – *D. litorellus*), ornamented mid legs are observed only in *D. pennatus* and *D. subpennatus*, while, on the contrary, the thickening of the costal vein is seen in *D. lineatocornis* and *D. litorellus*. Common characters for the species of this group are mostly black antennae, elongated postpedicel, yellow fore coxae, femora and tibiae, and dark hind coxae. Males of *D. lineatocornis* demonstrate moderate similarity with *D. pennatus* in wing shape, while the *D. subpennatus* wing shape is more similar to that of *D. argyrotarsis* (Fig. 3), which is also included in this species group by Pollet et al. (2010). Males of *D. lineatocornis* are similar to *D. argyrotarsis* based on morphometric leg characters (Fig. 4).
9. The group *Dolichopus plumitarsis* species group includes *D. simius, D. ptenopedilus* and *D. plumitarsis*. This group of species is characterized by the similar modification of the males fore leg – the fourth and fifth segments are black and enlarged. In addition, males have long cilia on the hind femur. From the point of view of leg morphometry, all three species of the group possess remarkable similarity (Fig. 4). In addition, a significant similarity in wing shape is noted between *D. plumitarsis* and *D. cilifemoratus*, whose males have modified fore legs, although a different type of modification (Fig. 3).
10. The group *Dolichopus ringdahli* species group includes *D. ringdahli* and *D. remipes*. In this group of species only *D. remipes* males has ornamented legs. Other than that, the species are similar in the following characters: completely black antennae, non-elongated postpedicel, a well-developed anal lobe, pale fore coxae and fore tibiae, and mostly pale hind tibia.

The groups of species identified by analyzing the morphometric features of the male’s wings, possessed, in most cases, similar limb modifications.

**Fig. 3.**
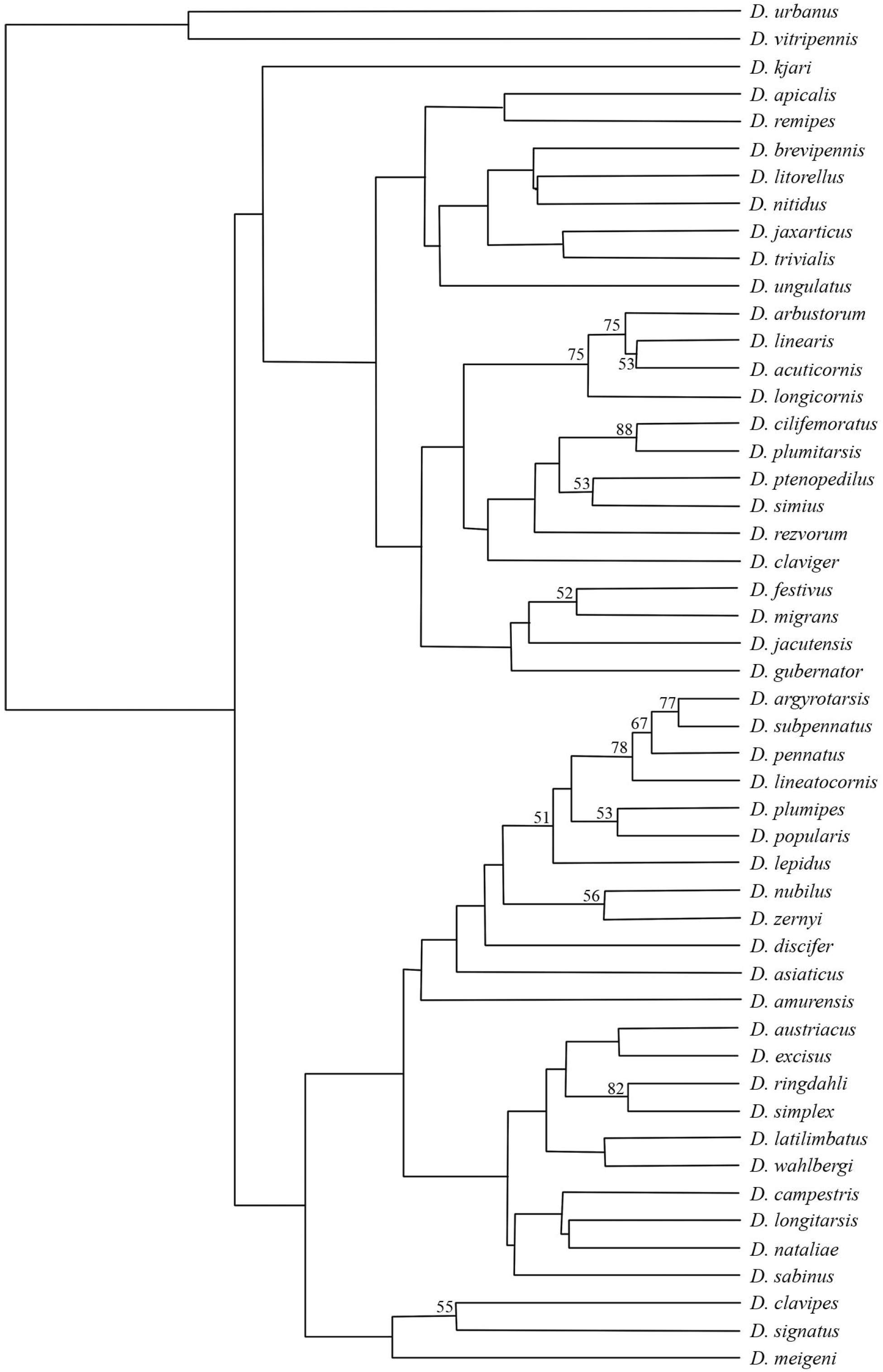
UPGMA tree from 1000 bootstrap replicates, obtained from 18 morphometric characters of wing shape. Bootstrap support values higher than 50% are indicated above branches.

**Fig. 4.**
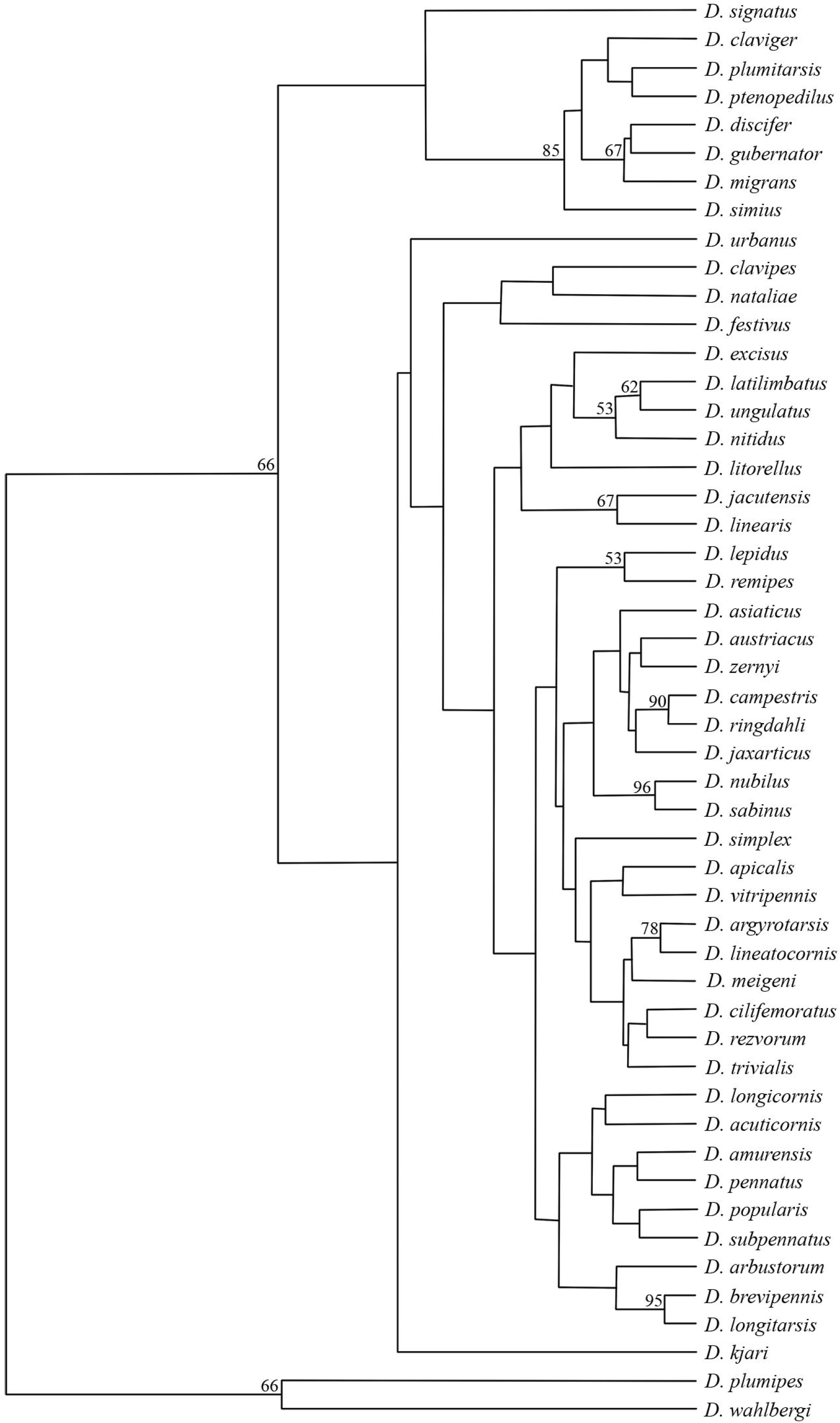
UPGMA tree from 1000 bootstrap replicates, obtained from 12 relative morphometric characters of legs. Bootstrap support values higher than 50% are indicated above branches.

Thus, the following groups were identified (Fig. 3):

1. *D. cilifemoratus* – *D. plumitarsis* (boot-strap support values, BS = 88) and *D. festivus* – *D. migrans* (BS = 52) – males of both species have foreleg modifications, only in males *D. cilifemoratus* and *D. festivus* – these are curved segments bearing protruding setae, and in *D. plumitarsis* and D. migrans – males have shortened and widened 4th and 5th segments of the fore tarsi.
2. *D. ptenopedilus* and *D. simius* male (BS = 53) have widened black 4th and 5th segments of the fore tarsi.
3. In the group *D. pennatus* – *D. subpennatus* – *D. argyrotarsis* – *D. lineatocornis* – *D. plumipes* – *D. popularis* (BS = 51) – only males of *D. lineatocornis* have no legs modifications, in other cases, the segments of the middle tarsi are widened and black or silver colored.

On the other hand, according to the wing shape, groups of species were identified that do not have significant similarity in molecular data and are not characterized by unique sets of morphological traits: *D. nubilus* – *D. zernyi* (BS = 56), *D. ringdahli* – *D. simplex* (BS = 82), *D. clavipes* – *D. signatus* (BS = 55), *D. acuticornis* – *D. longicornis* – *D. arbustorum* – *D. linearis* (BS = 75). Among the listed species, only *D. acuticornis* and *D. longicornis* are phylogenetically close (Fig. 2).

Identification of species groups based on the morphometric characters of legs is, in some cases, interrelated with the presence of modifications in the legs of males (Fig. 4). Thus, males of the species *D. discifer, D. gubernator, D. migrans, D. clavipes, D. plumitarsis, D. ptenopedilus* and *D. simius* (BS = 85) have a widened last segment of the fore legs, and males of *D. plumipes* and *D. wahlbergi* (BS = 66) have elongated and a feathered first segment of the middle tarsus.

The study of the characters phylogenetic signals showed that in each of the studied groups of characters (traditional morphological characters, characters of wing shape, and characters of legs morphometry) some carried a high phylogenetic signal, while, in a number of cases, the results were different for females and males.

Among the traditional morphometric characters, the most significant in the construction of a phylogenetic tree was shown: costal thickening at the tip of *R*_*1*_ (λ = 0.99, P <0.00001; K = 0.62, P = 0.005), the colour of the fore coxae (λ = 0.99, P <0.00001; K = 0.74, P = 0.007), the ornamented male’s foreleg (λ = 0.99, P <0.00001; K = 0.87, P <0.001), the colour of the femora (λ = 0.99, P <0.00001; K = 0.54, P = 0.03), the presence of cilia on the hind femora (λ = 0.99, P <0.00001; K = 0, 73, P <0.001), and the colour of the lower postocular cilia (λ = 0.99, P <0.00001; K = 0.59, P = 0.01).

Characters of wing shape also showed a high correlation with the phylogeny of the *Dolichopus* species. In males, the greatest value of phylogenetic signal value was demonstrated by the displacements of the following landmarks: X4 – terminus of *M*_*1+2*_ (λ = 0.80, P = 0.0005; K = 0.77, P = 0.003), X6 – the insertion point of *M*_*4*_ with *dm-m* (λ = 0.74, P = 0.0001; K = 0.52, P = 0.008), X7 – the insertion point of *M*_*1+2*_ and *dm-m* (λ = 0.74, P = 0.0006; K = 0, 58, P = 0.007), X8 and Y8 – the origin of *R*s (λ = 0.93, P <0.0001; K = 0.74, P <0.001 and λ = 0.55, P <0.01; K = 0.60, P = 0.002), Y9 – base of the anal vein (λ = 0.66, P = 0.005; K = 0.58, P = 0.005). The wing size of males had no significant phylogenetic signal.

In females, the greatest value of phylogenetic signal value was demonstrated by the displacements of the following landmarks: Y1 – terminus of *R*_*1*_ (λ = 0.99, P = 0.06; K = 0.67, P = 0.02), X3 – terminus of *R*_*4+5*_ (λ = 0.47, P = 0.04; K = 0.68, P = 0.009), X9 – base of the anal vein (λ = 0.99, P = 0.02; K = 0.75, P = 0.009). In addition, the wing size of females carried a significant phylogenetic signal: λ = 0.99, P <0.1; K = 0.68, P = 0.02. It should be noted that the relationship with phylogeny was mostly manifested by horizontal displacements of landmarks, which demonstrates a trend towards a wing length change.

Among the morphometric traits of males legs, the presence of a phylogenetic signal showed the following characters: F1/tar1 (λ = 0.90, P <0.0001; K = 0.48, P = 0.08), F2/T2 (λ = 0.44, P = 0.007; K = 0.44, P = 0.1). Among the morphometric leg traits of females: T1/tar1 (λ = 0.99, P = 0.02; K = 0.65, P = 0.006), F2/T2 (λ = 0.84, P = 0.037; K = 0, 64, P = 0.06), F3/tar3 (λ = 0.99, P = 0.02; K = 0.68, P = 0.03). Thus, it was shown that the most significant association with phylogeny in both males and females were characters of the limbs, such as the relative length of the first segment of the fore tarsus, which is usually associated with the presence of modifications in the fore tarsus of the male of the *Dolichopus* species.

## Discussion

The most commonly used morphological characters used for species and species-group diagnostics are the following: colour of legs, postocular cilia and squamal cilia, chaetotaxy of legs (the presence of a long apicoventral setae on the fore tibia of the male, the number of apical setae on the middle and hind femur, and the number of dorsal setae on the first segment of the hind legs), colour of antennae and various characters of sexual dimorphism. So, males often have an elongated postpedicel, long cilia on the hind femora, various legs ornament, and modified arista. Females of the *Dolichopus* species, sometimes not closely-related phylogenetically, are often morphologically similar, that causes difficulties in species diagnostics.

Interspecies differences in leg modifications, antennae, and other male secondary sexual characters suggest that sexual selection is a critical force influencing the evolution of the *Dolichopus* species. However, the main trends in the variability of these characters still remain unexplored, because a robust phylogenetic hypothesis should be based on a set of both morphological and molecular data. In this study, we carried out a comparative analysis of four different groups of characters, including both characters that are traditionally used for species diagnostics of *Dolichopus* species, and previously unexplored morphometric characters of wings and legs.

Molecular characters of 20 *Dolichopus* species, which have central- and eastern-Palaearctic habitats, were introduced into the analysis, and then the phylogeny of the genus was studied. When morphological and morphometric characters were superimposed on the constructed phylogenetic tree, the phylogenetic significance of each character was separately has been examined.

The greatest correlation with the phylogeny of the genus was demonstrated by male legs secondary sexual characters. Although various limb modifications are evident in males, females have also shown a significant phylogenetic signal of the same morphometric features as those in males.

This result suggests that complete separation between the sexes of the genetic mechanisms affecting legs modification is not necessary for the development of sexual dimorphism. In other words, females exhibit a similar, albeit less significant, change in the legs morphometric characteristics. Similar results, but for a different dimorphic structure – namely, eye stalks – were obtained for Diptera of the family Diopsidae (Baker & Wilkinson, 2001).

Reconstruction of characters of sexual dimorphism based on phylogenetic trees indicates that modifications of the males’ forelegs have evolved independently at least five times in the genus *Dolichopus*. Namely, separate groups such as *D. simius* – *D. ptenopedilus* – *D. plumitarsis, D. cilifemoratus* – *D. festivus* – *D. trivialis*, and in *D. migrans, D. gubernator, D. clavipes*. At least four times in the genus, modifications of the mid legs have evolved: in *D. pennatus – D. subpennatus*, separately for *D. argyrotarsis, D. popularis, D. plumipes*. In our study, there was only one species with hind leg modification, *D. remipes*, and it did not demonstrate a phylogenetic relationship with species with fore and middle leg modifications.

This result is preliminary, since many branches of the constructed phylogenetic trees do not have high bootstrap support, which means they may change when species will be added to the analysis. However, at this stage, it should be noted that the phylogenetically confirmed groups included species with modifications of the same pairs of legs: for example, there are mixed groups that include species with simple legs and species with modified legs, but never species with modification of the fore and middle legs.

Taking into account the high importance of flight in the life of the adult dolichopodid, the *Dolichopus* wing shape of has been repeatedly studied, and it was shown that the shape has a phylogenetic signal (Chursina & Negrobov, 2018); however, in this study, a more detailed examination of it was carried out separately for *Dolichopus* females and males. The analysis of the shape of the male wing allowed species with similar foot modifications to be classified into separate groups, which may mean an evolutionary relationship between the wing shape and the morphometric characters of the legs.

One of the possible explanations is that the wing shape is an adaptation to changes in the legs characters of males, but not females, since in females the phylogenetic signal shows other morphometric wing characters. Therefore, first, wing size did not have a significant phylogenetic signal in males, but it did in females. In addition, in males, such landmark displacements had a higher phylogenetic signal, which indicates a change in wing length and position of the posterior cross vein, which is most often associated with a change in the aerodynamic characteristics of flight (Weis-Fogh, 1973; Ellington, 1984), while in females, the leading edge of the wing remained the most phylogenetically significant. Landmarks described the shape of the wing base had a phylogenetic signal in both females and males.

In general, the results of this study show that evolutionary changes in the traits of sexual dimorphism correlate with changes in features such as legs morphometry and wing shape. Consequently, the evolutionary pattern of the *Dolichopus* species demonstrates a significant relationship between the variability of modifications in the limbs of males and the traits associated with it.

## Supporting information

Tble 2

## Supplementary Information

The online version contains supplementary material available at

## Acknowledgements

The work was funded by RFBR and NSFC according to the research project No 20-54-53005.

## Availability of data and material

The gene sequences were deposited in GenBank. The datasets generated during and/or analysed during the current study are available from the corresponding author on reasonable request.

## Author contribution

All authors contributed to the study conception and design. All authors read and approved the final manuscript.

## Declarations

### Ethics approval

Not applicable.

### Consent to participate

All authors have read and approved this manuscript.

### Consent for publication

All authors have read and approved this manuscript.

