## Supplementary material for "Phylogeny of the genus *Dolichopus* Latreille, 1796 and testing for phylogenetic signal in morphological characters and morphometric data": Table 1

**Table 1** The studied *Dolichopus* species, number of examined specimens, and GenBank accession number.

| № | Species | Number of specimens | | GenBank Accession No. |
|---|---|---|---|---|
| | | males | females | |
| 1 | *D. acuticornis* Wiedemann, 1817 | 14 | 36 | EU847538.1[A] |
| 2 | *D. amurensis* Stackelberg, 1930 | 5 | 2 | OK340616.1[*] |
| 3 | *D. apicalis* Zetterstedt, 1849 | 7 | 2 | OK491388.1[*] |
| 4 | *D. arbustorum* Stannius, 1831 | 24 | 20 | OK335810.1[*] |
| 5 | *D. argyrotarsis* Wahlberg, 1850 | 34 | 22 | OK335811.1[*] |
| 6 | *D. asiaticus* Negrobov, 1973 | 2 | 1 | OK340618.1[*] |
| 7 | *D. austriacus* Parent, 1927 | 17 | 9 | OK340619.1[*] |
| 8 | *D. brevipennis* Meigen, 1824 | 18 | 17 | AY744186.1[B] |
| 9 | *D. campestris* Meigen, 1824 | 19 | 39 | AY744212.1[B] |
| 10 | *D. cilifemoratus* Macquart, 1827 | 35 | 97 | AY958243.1[B] |
| 11 | *D. claviger* Stannius, 1831 | 20 | 14 | AY744206.1[B] |
| 12 | *D. clavipes* Haliday, 1832 | 2 | 1 | AY958248.1[B] |
| 13 | *D. discifer* Stannius, 1831 | 41 | 13 | AY744208.1[B] |
| 14 | *D. excisus* Loew, 1859 | 2 | 1 | AY958245.1[B] |
| 15 | *D. festivus* Haliday, 1832 | 1 | 2 | AY958236.1[B] |
| 16 | *D. gubernator* Mik, 1878 | 3 | 1 | OK446501.1[*] |
| 17 | *D. jacutensis* Stackelberg, 1929 | 5 | 2 | OK336092.1[*] |
| 18 | *D. jaxarticus* Stackelberg, 1927 | 8 | 3 | OK340613.1[*] |
| 19 | *D. kjari* Stackelberg, 1929 | 7 | 2 | OK340624.1[*] |
| 20 | *D. latilimbatus* Macquart, 1827 | 86 | 77 | AY744200.1[B] |
| 21 | *D. lepidus* Staeger, 1842 | 48 | 36 | AY744202.1[B] |
| 22 | *D. linearis* Meigen, 1824 | 19 | 28 | AY958239.1[B] |
| 23 | *D. lineatocornis* Zetterstedt, 1843 | 24 | 12 | OK340614.1[*] |
| 24 | *D. litorellus* Zetterstedt, 1852 | 4 | 2 | OK340622.1[*] |
| 25 | *D. longicornis* Stannius, 1831 | 82 | 36 | AY958240.1[B] |
| 26 | *D. longitarsis* Stannius, 1831 | 95 | 110 | OK336131.1[*] |
| 27 | *D. meigeni* Loew, 1857 | 12 | 3 | OK491386.1[*] |
| 28 | *D. migrans* Zetterstedt, 1843 | 36 | 30 | OK446551.1[*] |
| 29 | *D. nataliae* Stackelberg, 1930 | 4 | 3 | OK340621.1[*] |
| 30 | *D. nitidus* Fallen, 1823 | 10 | 6 | EU847539.1[A] |
| 31 | *D. nubilus* Meigen, 1824 | 4 | 1 | AY958244.1[B] |
| 32 | *D. pennatus* Meigen, 1824 | 41 | 40 | OK446503.1[*] |
| 33 | *D. plumipes* (Scopoli, 1763) | 46 | 44 | EU847548.1[A] |
| 34 | *D. plumitarsis* Fallen, 1823 | 18 | 2 | OK340623.1[*] |
| 35 | *D. popularis* Wiedemann, 1817 | 9 | 19 | AY744190.1[B] |
| 36 | *D. ptenopedilus* Meuffels, 1982 | 10 | 2 | OK340617.1[*] |
| 37 | *D. remipes* Wahlberg, 1839 | 13 | 27 | OK446520.1[*] |
| 38 | *D. rezvorum* Stackelberg, 1930 | 19 | 3 | OK489436.1[*] |
| 39 | *D. ringdahli* Stackelberg, 1930 | 74 | 49 | OK491385.1[*] |
| 40 | *D. sabinus* Haliday, 1838 | 9 | 4 | OK336364.1[*] |
| 41 | *D. signatus* Meigen, 1824 | 2 | 1 | AY958235.1[B] |
| 42 | *D. simius* Parent, 1927 | 6 | 2 | OK340615.1[*] |
| 43 | *D. simplex* Meigen, 1824 | 41 | 42 | AY744203.1[B] |
| 44 | *D. subpennatus* d'Assis Fonseca, 1976 | 4 | 6 | OK446507.1[*] |
| 45 | *D. trivialis* Haliday, 1832 | 2 | 5 | AY744210.1[B] |
| 46 | *D. ungulatus* (Linnaeus, 1758) | 201 | 112 | EU847559.1[A] |

| | | | | |
|---|---|---|---|---|
| 47 | *D. urbanus* Meigen, 1824 | 1 | 1 | AY744182.1[B] |
| 48 | *D. vitripennis* Meigen, 1824 | 1 | 1 | AY744195.1[B] |
| 49 | *D. wahlbergi* Zetterstedt, 1843 | 1 | 2 | EU847561.1[A] |
| 50 | *D. zernyi* Parent, 1927 | 11 | 8 | OK340620.1[*] |

[A]Germann et al. (2010). [B]Bernasconi et al. (2007b). [*]sequence have been obtained by the authors of this study and deposited in GenBank.
