## Supplementary material for "Phylogeny of the genus *Dolichopus* Latreille, 1796 and testing for phylogenetic signal in morphological characters and morphometric data": Tble 2

**Table 2** Overview of phylogenetic analysis results of 50 *Dolichopus* species for selected nodes.

| *Dolichopus* species groups | MP | ML | ME | NJ |
|---|---|---|---|---|
| *D. acuticornis – D. longicornis* | 79 | 83 | 93 | 89 |
| *D. arbustorum – D. argyrotarsis* | 100 | 100 | 100 | 100 |
| *D. asiaticus – D. zernyi* | 92 | 96 | 99 | 99 |
| *D. arbustorum – D. argyrotarsis – D. zernyi – D. asiaticus* | 86 | 95 | 90 | 97 |
| *D. cilifemoratus – D. festivus – D. trivialis* | 97 | 99 | 99 | 99 |
| *D. kjari – D. jacutensis* | 100 | 100 | 100 | 100 |
| *D. nubilus – D. latelimbatus – D. excisus* | 99 | 99 | 99 | 99 |
| *D. plumipes – D. simplex* | 100 | 100 | 100 | 100 |
| *D. pennatus – D. subpennatus – D. lineatocornis– D. litorellus* | 85 | 88 | 75 | 82 |
| *D. popularis – D. urbanus* | 100 | 99 | 100 | 100 |
| *D. ringdahli – D. remipes* | 94 | 90 | 99 | 98 |
| *D. simius – D. ptenopedilus – D. plumitarsis* | 100 | 100 | 100 | 100 |
